# Transient Overexpression of VEGF-A in Adipose Tissue Promotes Energy Expenditure via Activation of the Sympathetic Nervous System

**DOI:** 10.1101/323659

**Authors:** Yueshui Zhao, Xin Li, Li Yanga, Kristin Eckel-Mahan, Qingchun Tong, Xue Gu, Mikhail G. Kolonin, Sun Kai

## Abstract

Adipose-derived VEGF-A stimulates functional blood vessel formation in obese fat pads which in turn facilitates healthy expansion of the adipose tissue. However, the detailed mechanism(s) governing the process remains largely unknown. Here, we investigated the role of sympathetic nervous system activation in the process. To this end, we induced overexpression of VEGF-A in an adipose-specific doxycycline (Dox)-inducible transgenic mouse model for a short period of time during high fat-diet (HFD) feeding. We found that local overexpression of VEGF-A in adipose tissue stimulated lipolysis and browning rapidly after Dox induction. Immunofluorescent staining against tyrosine hydroxylase (TH) indicated higher levels of sympathetic innervation in adipose tissue of transgenic mice. In response to the increased norepinephrine (NE) level, expression of β3-andrenoceptor was significantly upregulated and the downstream protein kinase A (PKA) pathway was activated, as indicated by the enhanced phosphorylation of the whole PKA substrates, in particular the hormone sensitive lipase (HSL) in adipocytes. As the result, the adipose tissue exhibited increased lipolysis, browning, and energy expenditure. Importantly, all these effects were abolished upon the treatment with β3-adrenoceptor antagonist SR59230A. Collectively, these results demonstrate that transient overexpressed VEGF-A activates sympathetic nervous system which hence promotes lipolysis and browning in adipose tissue.

## Introduction

Obesity, characterized by an excess of white adipose tissue (WAT), is a major risk factor for many chronic disorders, including type 2 diabetes, cardiovascular diseases (CVD) and hypertension [1–3]. Adipose tissue has the ability for lifelong growth and almost unlimited expansion. Adipocyte differentiation and adipose tissue growth rely on the formation of new blood vessels that support proper oxygen, nutrition and stem and precursor cells [3]. However, during obesity development, the synchrony between adipose tissue expansion and vascularization is lost, giving rise to a local hypoxic condition [1, 4–6], which further results in fibrosis, inflammation and systemic metabolic disorders, such as insulin resistance and atherosclerosis [4, 5, 7, 8]. Hence, improving the vascular network and reversing hypoxia in the pathologically expanded WAT by angiogenesis have been proposed to be a potential strategy to deal with obesity and obesity-related metabolic disorders [2–4, 6].

Multiple angiogenic factors, such as vascular endothelial, hepatocyte, or fibroblast growth factors (VEGF, HGF, FGF) as well as angiopoietin-1 (Ang-1) and Ang-2 are secreted by adipocytes, suggesting that they auto-regulate vascular homeostasis [3]. Among them, VEGF-A is the only *bona fide* endothelial cell growth factor that has been found to account for most of the proangiogenic activity in adipose tissue [9, 10]. VEGF-A levels in adipose tissue are significantly upregulated after exercise and cold exposure [11, 12]. However, adipose tissue frequently fails to mount high enough levels of VEGF-A in response to hypoxia in obese adipose tissue, which leads to lack of angiogenesis in obese adipose tissue [5]. Interestingly, we recently applied a doxycycline (Dox)-inducible mouse model that specifically overexpresses VEGF-A in adipose tissue and found that locally overexpressed VEGF-A stimulated functional new blood vessels by angiogenesis. We further demonstrated that long-term angiogenesis facilitated adipose tissue expansion by reducing the local hypoxia, fibrosis and inflammation [13, 14]. As a result, the mice remained metabolically fit under HFD, with enhanced energy expenditure and improved insulin sensitivity [13, 14]. Of note, among all the metabolically beneficial effects of VEGF-A in obese adipose tissue, the most significant one is a higher rate of energy expenditure [14, 15]. As the result, the VEGF-A transgenic (Tg) mice gained less body weight despite unchanged food intake [15]. Importantly, we previously found that under HFD challenge, overexpression of VEGF-A resulted in features characteristic of brown (beige) adipocytes in sWAT of VEGF-A Tg mice [14]. Confirming sWAT browning, we observed elevated expression levels of brown-fat-associated proteins, including uncoupling protein-1 (UCP-1) and PPARγ coactivator-1α (PGC-1α) in subcutaneous WAT (sWAT) [14]. These findings were confirmed subsequently by other reports [11, 15, 16].

Because previous studies have focused on long-term VEGF-A effects, it has remained unclear if angiogenesis was responsible for WAT browning or whether VEGF-A has direct effects on other cells. To investigate short-term VEGF-A signaling in WAT, we took advantage of the Dox-inducible adipocyte-specific VEGF-A transgenic mouse model that we previously reported [14]. To determine the direct function of VEGF-A and rule out the possible secondary effects during chronic obesity, in the current study we induced the VEGF-A expression by Dox for only a short period of time (one week) under HFD challenging. Surprisingly, even after the short-term induction, VEGF-A triggered denser local sympathetic nerve fiber innervation which further stimulated the upregulation of β3-adrenoceptor and the activation of downstream PKA signaling pathway, ultimately leading to increased lipolysis and browning in adipose tissue. Thus our results provide a greater understanding of the molecular mechanism by which VEGF-A stimulates adipose tissue activity to accelerate energy expenditure.

## Results

### Transient overexpression of VEGF-A in adipose tissue reduced fat mass and enhanced energy expenditure after HFD feeding

Previously we reported that adipose-derived VEGF-A in chronically HFD-challenged mice caused dramatically reduced fat mass size and body weight, which was associated with improved insulin sensitivity, as well as enhanced energy expenditure [17]. Here, we challenged the VEGF-A Tg mice with HFD plus 30 mg/kg Dox for only 7 days to rule out the potential secondary metabolic effects of chronic diet-induced obesity. To our surprise, even after the short term induction, VEGF-A Tg mice gained less body weights and had smaller fat pads compared to their littermate controls (**Fig. 1A-1C**). Moreover, histological examination revealed a smaller adipocyte size in WAT of VEGF-A Tg mice (**Fig. 1D**).

**Figure 1.**
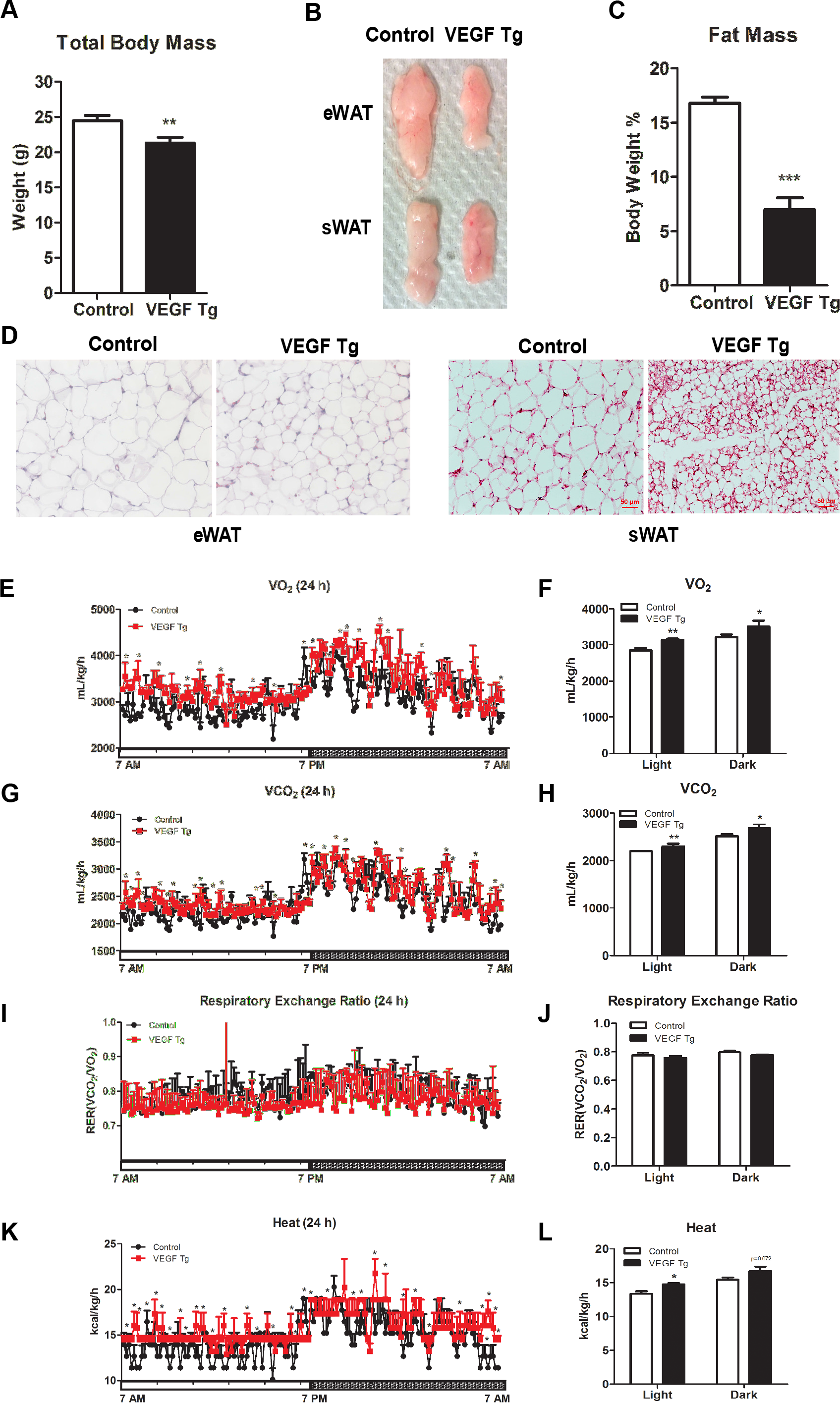
VEGF-A transgenic mice exhibited smaller adipocytes and reduced total fat mass shortly after VEGF-A induction. **(A)** Body weights of VEGF Tg mice and their littermate controls after HFD (60% fat) plus 30 mg/kg Dox feeding for 7 days (n = 6 per group, Student’s *t-test*, **, *p* < 0.01). **(B)** Comparison of the sizes of different adipose tissues (eWAT and sWAT) collected from VEGF Tg mice and their littermate controls after HFD plus Dox feeding for 7 days. **(C)** MRI analysis of fat mass in VEGF Tg mice and their littermate controls after HFD plus Dox feeding for 7 days (n = 6 per group, Student’s *t-test*, ***, *p* < 0.005). **(D)** H & E staining of different adipose tissue sections (eWAT and sWAT) collected from VEGF Tg mice and their littermate controls after HFD plus Dox feeding for 7 days (Scale bar, 50 μm). (**E-L**) Indirect calorimetry performed in a CLAMS system after HFD plus Dox feeding for 7 days. (**E**) O_2_ consumption profile of VEGF-A Tg and control mice during a 12 h-12 h light-dark cycle (VO_2_). (**F**) Histogram representative of light time and dark time of result in (E).(**G**) CO2 production (VCO2) profile of VEGF-A Tg and control mice during a 12 h-12 h light-dark cycle. (**H**) Histogram representative of light time and dark time of result in (G). (**I**) RER (VCO_2_/VO_2_) profile of VEGF-A Tg and control mice during a 12 h-12 h light-dark cycle. (**J**) Histogram representative of light time and dark time of result in (G) (n = 3 per group, Student’s *t-test*, **, *p* < 0.01; *, *p* < 0.05). **(K)** Heat generation profile of VEGF-A Tg and control mice during a 12 h-12 h light-dark cycle. (**J**) Histogram representative of light time and dark time of result in (K) (n = 3 per group, Student’s *t-test*, *, *p* < 0.05).

Indirect calorimetry indicated that oxygen consumption (VO_2_, **Fig. 1E** **and 1F**) and CO2 production (VCO_2_, **Fig. 1G** **and 1H**) were significantly increased in VEGF-A Tg mice, indicating higher energy turnover in these mice. The respiratory exchange ratio (RER, VCO_2_/VO_2_) showed no difference between VEGF-A Tg mice and their littermate controls, indicating no change in glucose and lipid turnover between the two groups (**Fig. 1I** **and 1J**). Importantly, heat generation was significantly increased in the VEGF-A transgenic mice (**Fig. 1K** **and 1L**), suggesting that overexpressed VEGF-A in adipose tissue increases thermogenesis.

The results suggested that short-term induction of VEGF-A in adipose tissue leads to less body weights, smaller fat mass and a higher rate of energy expenditure, suggesting a direct local role of adipose VEGF-A in whole body metabolism.

### Local overexpression of VEGF-A in adipose tissue increased mitochondrial biogenesis and function

Mitochondria is the predominant organelle for energy production in adipose tissues [18]. To determine whether mitochondrial number and function are regulated by VEGF-A in adipose tissue, we first measured the β-oxidation related gene expression and found that most of this group of genes were significantly upregulated in sWAT of the VEGF-A Tg mice (**Fig. 2A**). We then examined the effect of VEGF-A overexpression on mitochondrial biogenesis. Analysis of mitochondrial biogenesis gene expression revealed that *Pgc1α, Nrf1* and *Cox IV* were all upregulated in the sWAT of the VEGF-A Tg mice (**Fig. 2B**). The levels of mitochondrial DNA (so called mtDNA) were significantly higher (**Fig. 2C**), further supporting the increase in mitochondrial content in the sWAT of the VEGF-A Tg mice. In line with the qPCR and mtDNA results, Immunofluorescent staining with an antibody to COX IV, the biomarker of mitochondria, showed that the positive green fluorescent signals were much stronger in the sWAT of the VEGF-A Tg mice. Furthermore, Western blotting using an antibody against SDHA, another mitochondrial protein indicated more SDHA proteins in the sWAT of the VEGF-A Tg mice.

**Figure 2.**
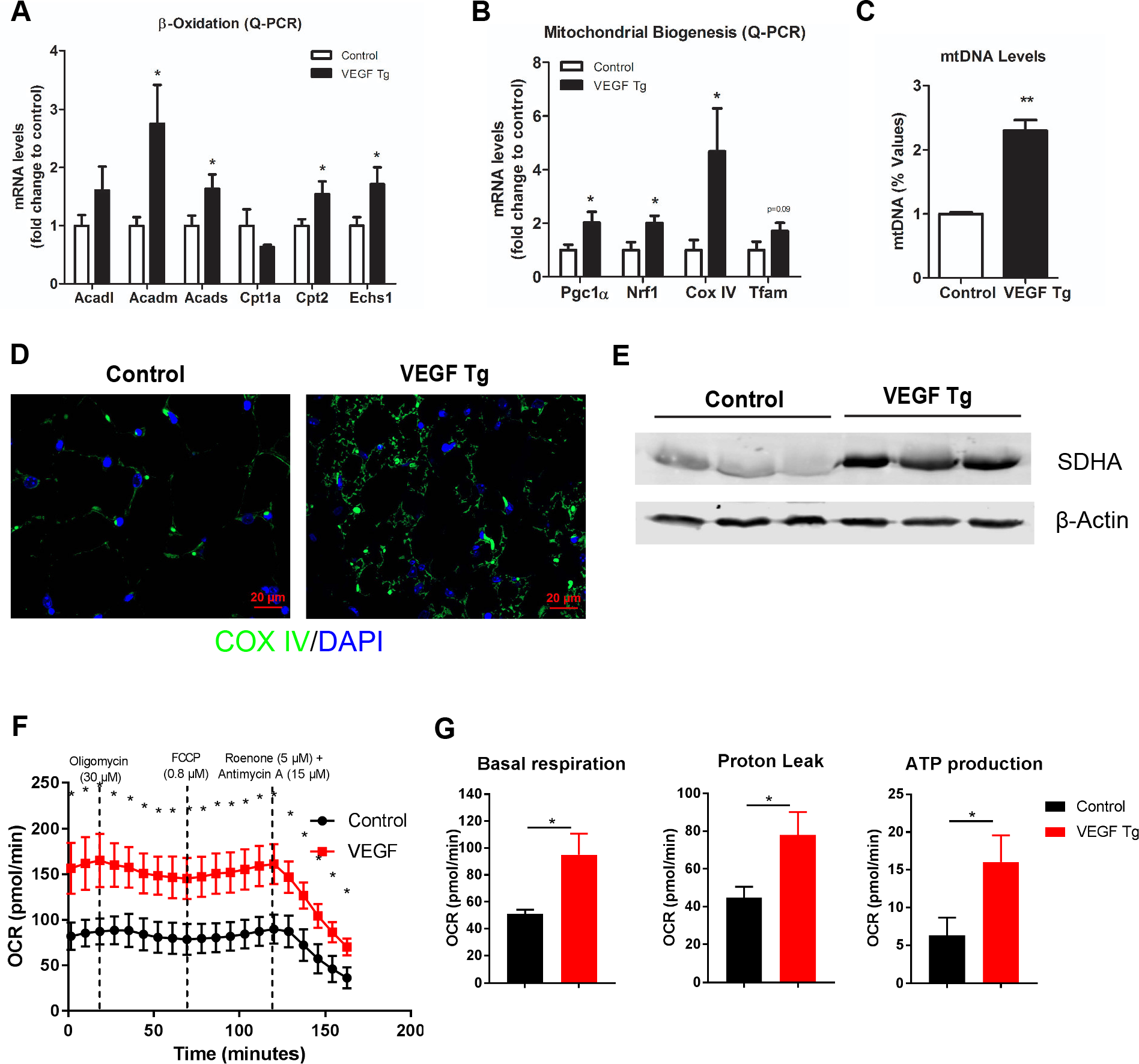
Local overexpression of VEGF-A stimulates mitochondrial biogenesis and functions in AT. **(A)** Q-PCR analysis of β-oxidation-related genes, namely *Acadl, Acadm, Acads, Cptla, Cpt2*, and *Echsl* in sWAT of VEGF Tg mice and their littermate controls after HFD plus Dox feeding for 7 days (n = 5 per group, Student’s *t-test*, *, *p* < 0.05). **(B)** Q-PCR analysis of mitochondrial biogenetic genes including *Pgcla, Nrf1, Cox IV*, and *Tfam* in sWAT of VEGF Tg mice and their littermate controls after HFD plus Dox feeding for 7 days (n = 5 per group, *Student’s t-test*, *, *p* < 0.05). **(C)** The mitochondrial DNA (mtDNA) content in sWAT of VEGF Tg mice and their littermate controls after HFD plus Dox feeding for 7 days. The copy number of mtDNA was calculated by the ratio of the mtDNA gene NADH dehydrogenase alpha 1 (*Nadha1*) to the nuclear gene lipoprotein lipase (*Lpl*) (n = 5 per group, *Student’s t-test*, **, *p* < 0.01). **(D)** Immunofluorescent (IF) staining with α-COX IV antibody (green) in sWAT of VEGF Tg mice and their littermate controls after HFD plus Dox feeding for 7 days. The nuclei were stained with DAPI (blue) (Scale bar, 20 μm). **(E)** Western blotting analysis of protein levels of SDHA with α-SDHA antibody in sWAT of VEGF Tg mice and their littermate controls after HFD plus Dox feeding for 7 days. Equal loading control was demonstrated by α-β-actin antibody (n = 3 per group. *Student’s t-test*, *, *p* < 0.05, **, *p* < 0.01). **(F)** Oxygen consumption rate (OCR) in sWAT measured by seahorse machine. The sWAT was collected from VEGF Tg and their littermate control mice after 7 days DOX induction. The injection time and final concentrations of the different compounds used for the assay were shown at the above of the curves. Results were represented as mean° S.E.M., n=3 per group. *Student’s t-test*, *, *p* < 0.05). **(G)** Bioenergetic parameters inferred from OCR traces. Results were represented as mean° S.E.M., n=3 per group. Student’s *t-test*, *, *p* < 0.05, ***, p<0.001).

We then measured the function of mitochondria in the adipose tissue of the VEGF-A Tg mice by Seahorse. Fat pads collected from VEGF-A Tg mice consistently exhibited more oxygen consumption rate (OCR) at different time windows upon different compound treatment when compared to the controls (**Fig. 2F**). Detailed analyses at the basal level, during proton leak and ATP generation stages all indicated more OCR in the WATs of the transgenic mice (**Fig. 2G**). Our results clearly show that mitochondrial content and function were increased in sWAT of the VEGF-A Tg mice.

### Local overexpression of VEGF-A in adipose tissue results in enhanced lipolysis

Previously we and others reported that VEGF-A promotes adipocyte browning, manifested as multilocular lipid droplets in sWAT [11, 14, 19]. Here, we further found that in addition to the multilocular structures, lipid droplets in WAT of the transgenic mice were smaller in size (**Fig. 1D**). Since the balance between lipid synthesis (lipogenesis) and lipid breakdown (lipolysis) determines the total lipid levels in adipocytes [20], we next analyzed the expression levels of key enzymes involved in these processes. The qPCR results reveal that the levels of lipogenic enzymes, i.e. acetyl-CoA carboxylase 1 (*Acc1*) and fatty acid synthase (*Fasn*) were dramatically downregulated (**Fig. 3A**), while the levels of the lipolytic enzyme adipose triglyceride lipase (*Atgl*), hormone-sensitive lipase (*Hsl*) and lipase cofactor α/β hydrolase domain containing protein 5 (*Abhd5*, also known as CGI-58) were significantly upregulated in the sWAT of VEGF-A Tg mice (**Fig. 3B**). Of note, the mRNA levels of CD36, the key fatty acid transporter in adipose tissue [21], were also significantly upregulated in the sWAT of VEGF-A Tg mice (**Fig. 3C**).

**Figure 3.**
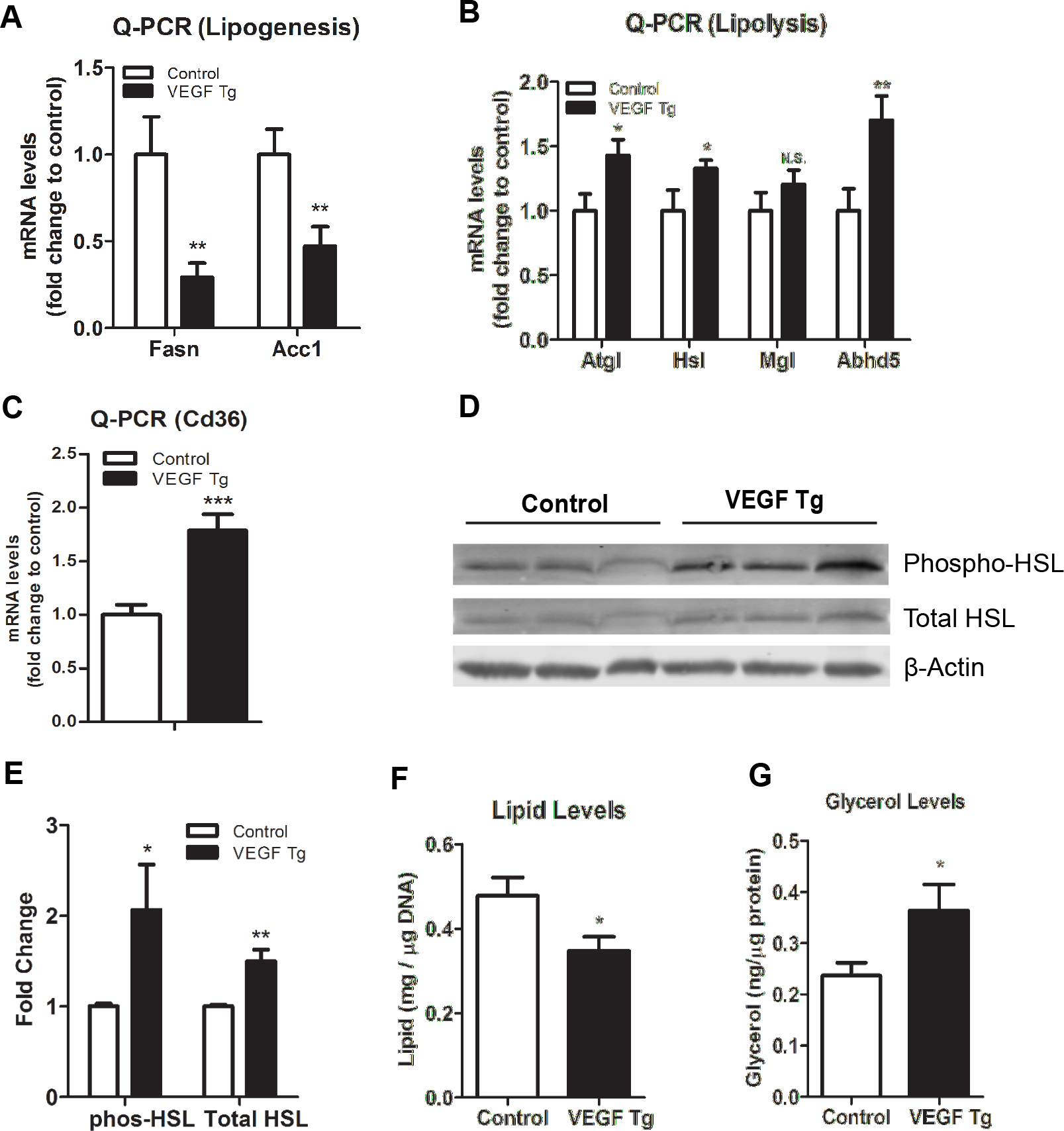
Local overexpression of VEGF-A stimulates lipolysis in WAT. **(A)** Q-PCR analysis of lipogenic genes, namely *Acc1* and *Fasn* in sWAT of VEGF Tg mice and their littermate controls after HFD plus Dox feeding for 7 days (n = 6 per group, *Student’s t-test*, **, *p* < 0.01). **(B)** Q-PCR analysis of lipolytic genes, namely *Atgl, Hsl, Mgl* and *Abhd5* in sWAT of VEGF Tg mice and their littermate controls after HFD plus Dox feeding for 7 days (n = 6 per group, *Student’s t-test*, *, *p* < 0.05, **, *p* < 0.01, n.s.: no statistical significance). **(C)** Q-PCR analysis of *Cd36* gene in sWAT of VEGF Tg mice and their littermate controls after HFD plus Dox feeding for 7 days (n = 6 per group, *Student’s t-test*, ***, *p* < 0.005). **(D and E)** Western blotting (D) and quantification of the band density on the Western blots by ImageJ software (E) for phosphorylated HSL at serine 660 (P^ser660^) as well as total HSL levels in sWAT of VEGF Tg mice and their littermate controls after HFD plus Dox feeding for 7 days. Results were normalized by P-actin (n = 3 per group. *Students t-test*, *, *p* < 0.05, **, *p* < 0.01). **(F)** Total triglyceride content in sWAT of VEGF Tg mice and their littermate controls after HFD plus Dox feeding for 7 days. (n = 4~5 per group. *Students t-test*, *, *p* < 0.05). **(G)** Glycerol levels (as the product of lipolysis) in sWAT of VEGF Tg mice and their littermate controls after HFD plus Dox feeding for 7 days (n = 5 per group. *Student’s t-test*, *, *p* < 0.05).

In line with the upregulation of mRNA levels of the lipolytic genes, Western Blotting results indicated that the total protein levels of HSL, an enzyme rate-limiting lipolysis [22], were increased in the sWAT of VEGF-A Tg mice. Importantly, the levels of phosphorylated HSL (a marker of its activity) were also significantly increased (**Fig. 3D**, quantitative measurement in **Fig. 3E**), suggesting the function of HSL was enhanced in the WAT of the VEGF-A Tg mice. Consequently, we observed that the total triglyceride levels were decreased, while the levels of glycerol, a product of the lipolytic process, were dramatically increased (**Fig. 3F** **and 3G**). Collectively, our results suggest that adipose-derived VEGF-A triggers enhanced lipolysis in WAT.

### Local overexpression of VEGF-A in adipose tissue stimulated sympathetic neuronal innervation and activation

Next, we sought to investigate the mechanism(s) governing the browning and lipolytic effects induced by VEGF-A. Since sympathetic activation plays a critical role in browning and lipolysis in WAT [23–25], we thus determined whether overexpression of VEGF-A in adipose tissue stimulates local sympathetic activation. To this end, we first examined the nerve density by whole mount immunofluorescent staining with tyrosine hydroxylase (TH), the marker of the sympathetic neurons in sWAT of the VEGF-A Tg and their littermate control mice. Interestingly, we found a dramatic increase in the density of the sympathetic nerve fibers in the sWAT of VEGF-A Tg mice (**Fig. 4A**), suggesting an enhanced innervation in these adipose tissues. We then analyzed several markers for sympathetic nervous system and found that the levels of norepinephrine (NE), the marker of innervated sympathetic neurons [23], were significantly higher in sWAT of the VEGF-A Tg mice (**Fig. 4B**, **left**). Importantly, the serum NE levels did not change between the groups (**Fig. 4B**, **right**), indicating the local innervation effect of VEGF-A.

**Figure 4.**
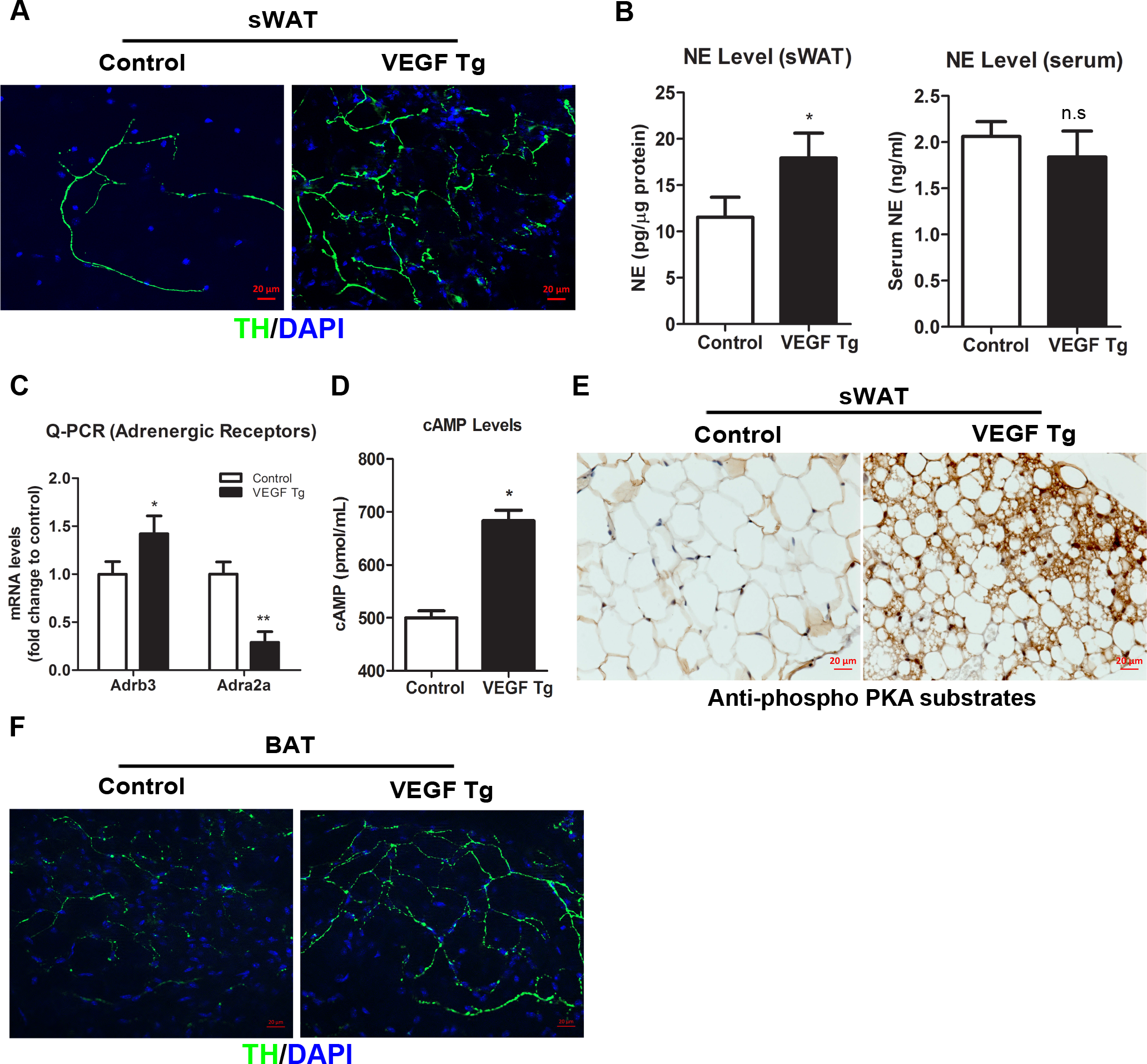
Local overexpression of VEGF-A induces sympathetic activation in WAT. **(A)** Whole mount IF staining with α-tyrosine hydroxylase (TH) antibody (green) in sWAT of VEGF Tg mice and their littermate controls after HFD plus Dox feeding for 7 days. The nuclei were stained with DAPI (blue) (representative of 3 trials. Scale bar, 20 μm). **(B)** NE levels in sWAT (left) and serum (right) of VEGF Tg mice and their littermate controls after HFD plus Dox feeding for 7 days. Results for sWAT were normalized with the total protein levels (n = 5 for control group, n=6 for VEGF Tg group. *Student’s t-test*, *, p < 0.05, n.s: no statistical significance). **(C)** Q-PCR analysis of adrenergic receptor (AR) genes, namely *Adrb3* and *Adra2a*, in sWAT of VEGF Tg mice and their littermate controls after HFD plus Dox feeding for 7 days (n = 6 per group, *Student’s t-test*, *, *p* < 0.05; **, *p* < 0.01). **(D)** cAMP levels in sWAT of VEGF Tg mice and their littermate controls after HFD plus Dox feeding for 7 days (n = 5 per group. *Students t-test*, *, *p* < 0.05). **(E)** IHC staining with α-total phosphorylated PKA substrate antibody in sWAT of VEGF Tg mice and their littermate controls after HFD plus Dox feeding for 7 days (Scale bar, 100 μm). **(F)** Whole mount IF staining with α-tyrosine hydroxylase (TH) antibody (green) in BAT of VEGF Tg mice and their littermate controls after HFD plus Dox feeding for 7 days. The nuclei were stained with DAPI (blue) (representative of 3 trials, scale bar, 20 μm).

Since NE triggers the downstream signaling events via β3-adrenergic receptor (β3-AR, *Adrb3*) pathway [26], we next determined the expression levels of β3-AR and its counteracting partner, Adra2α in the sWAT of VEGF-A Tg mice. As anticipated, the qPCR data indicated that the mRNA levels of β3-AR was significantly increased, whereas the levels of Adra2α was dramatically decreased in the sWAT of VEGF-A Tg mice (**Fig. 4C**). In line with the upregulation of β3-AR levels, the levels of cAMP as well as the levels of phosphorylated PKA substrates were also dramatically increased in sWAT of VEGF-A Tg mice (**Fig. 4D&E**), indicating the activation of protein kinase A (PKA) pathway, the classical downstream event of stimulated sympathetic tone [27].

Since BAT is another metabolically active adipose tissue [25], we decided to further test sympathetic innervation in this fat pad. As anticipated, the density of the sympathetic nervous system was also enhanced in the BAT of VEGF-A Tg mice (**Fig. 4F**).

Taken together, our results suggested that the sympathetic tone and its downstream β3-adrenergic signaling pathway are activated by VEGF-A in the sWAT and BAT of the transgenic mice.

### Adipose VEGF-A-induced metabolic effects were suppressed by β3-AR antagonist SR59230A

To further confirm the critical role of β3-AR signaling in VEGF-A mediated metabolic improvements, we blocked the pathway by administrating the selective β3-AR antagonist SR59230A into the VEGF-A Tg and the control mice by intraperitoneal injection (i.p.). We found that after i.p. injection of SR59230A, cAMP levels induced by VEGF-A were dramatically reduced (**Fig. 5A**). To further determine the effect of SR59230A on PKA activity, we measured the total PKA phosphorylated substrate proteins by IHC and we found the signal of the positive stains were dramatically suppressed upon SR59230A treatment (**Fig. 5B**, compare the right to the middle panel). Histologically, we found that blockade of β3-AR by SR59230A enlarged the fat mass and the adipocyte size in the VEGF-A Tg mice (**Fig. 5C** **and 5D**). As anticipated, the lipolytic effect induced by VEGF-A was impaired by blockade of β3-AR, as shown by the downregulation of the critical lipolytic enzymes at both the mRNA and protein levels (**Fig. 5E&F**, quantitative measurement in **Fig. 5G**). Consequently, Glycerol production induced by VEGF-A in sWAT was dramatically reduced after SR59230A treatment (**Fig. 5H**). Furthermore, the IHC staining using an antibody to UCP1 showed significantly decreased UCP1 in sWAT of the VEGF-A Tg mice upon SR59230A treatment compared to controls (**Fig. 5I**, compare the right to the middle panel), suggesting that the browning effect of VEGF-A was abolished by the blockage of β3-AR in the sWAT. Collectively, our results suggest that the β3-AR signaling pathway is necessary and sufficient for the VEGF-A-induced metabolic effects in adipose tissue.

**Figure 5.**
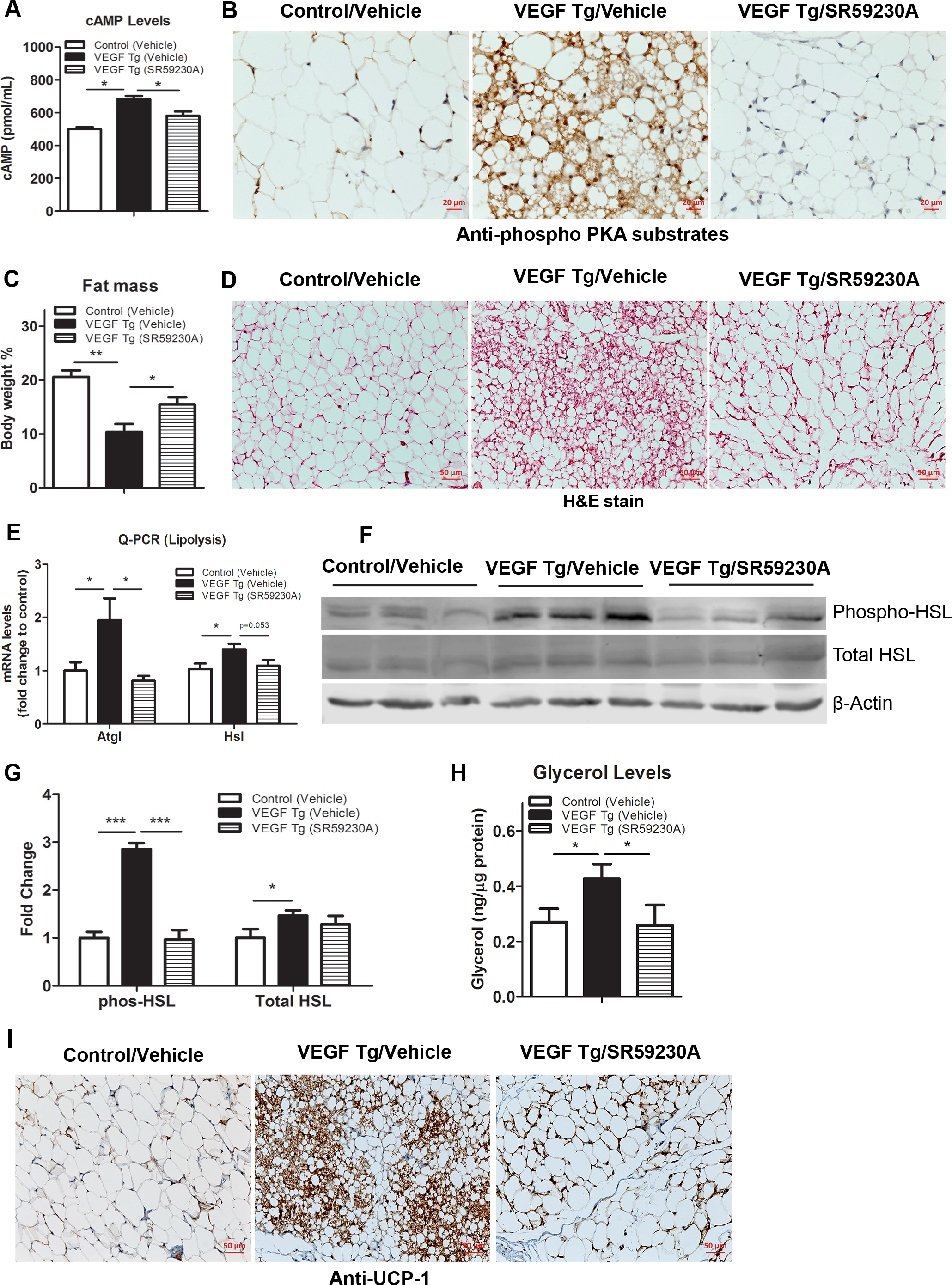
SR59230A, a selective β3-AR antagonist, suppresses VEGF-A-induced lipolysis and browning effect in WAT. **(A)** cAMP levels in sWAT of VEGF Tg mice and their littermate controls after HFD plus Dox feeding together with or without SR59230A treatment for 7days (n = 4 for control group, and n=5 for VEGF-A Tg group. Student’s *t-test*, *, *p* < 0.05). **(B)** IHC staining with α-total phosphorylated PKA substrate antibody in sWAT of VEGF Tg mice and their littermate controls after HFD plus Dox feeding together with or without SR59230A treatment for 7 days (Scale bar, 20 μm). **(C)** MRI analysis of fat mass in VEGF Tg mice and their littermate controls after HFD plus Dox feeding at the presence of either vehicle (1% DMSO in PBS) or SR59230A (via i.p. injection) for 7 days. Fat mass were normalized with total body mass (n = 6 per group, Student’s *t-test*, *, *p* < 0.05; **, *p* < 0.01). **(D)** H & E staining of sWAT collected from VEGF Tg mice and their littermate controls after HFD plus Dox feeding at the presence or absence of SR59230A for 7 days (Scale bar, 50 μm). **(E)** Q-PCR analysis of lipolytic genes, namely *Atgl* and *Hsl*, in sWAT of VEGF Tg mice and their littermate controls after HFD plus Dox feeding at the presence or absence of SR59230A for 7 days (n = 5 per group, Student’s *t-test*, *, *p* < 0.05). **(F and G)** Western blotting (F) and quantitative measurement of band’s density on the Western blots by ImageJ software (G) for phosphorylated HSL at serine 569/660 (P^ser569/660^) as well as total HSL levels in sWAT of VEGF Tg mice and their littermate controls after HFD plus Dox feeding at the presence or absence of SR59230A for 7days. Results were normalized with β-actin (n = 3 per group. Student’s *t-test*, *, *p* < 0.05, ***, *p* < 0.005). **(H)** Glycerol levels (product of lipolysis) in sWAT of VEGF Tg mice and their littermate controls after HFD plus Dox feeding at the presence or absence of SR59230A for 7 days (n = 4 for control group, and n=5 for VEGF-A Tg group. Student’s *t-test*, *, *p* < 0.05). **(I)** IHC staining with α-UCP1 antibody in sWAT of VEGF Tg mice and their littermate controls after HFD plus Dox feeding together with or without SR59230A for 7 days (Scale bar, 50 μm).

Based on these results, we present a working model (**Fig. 6**). According to the model, adipose-derived VEGF-A stimulates sympathetic innervation in adipose tissue. NE, locally released by the stimulated sympathetic nerve fibers, triggers the activation of β3-AR signaling which in turn induces higher levels of cAMP resulting in activation of PKA. The activated PKA phosphorylates the rate-limiting lipolytic enzyme HSL and promotes lipolysis in adipocytes; Furthermore, PKA phosphorylates critical transcriptional factors which directly upregulate the expression of UCP1 and browning in sWAT [28, 29]. Both lipolysis and browning facilitate the lipid breakdown as well as heat generation in mitochondria which ultimately result in enhanced energy expenditure.

**Figure 6.**
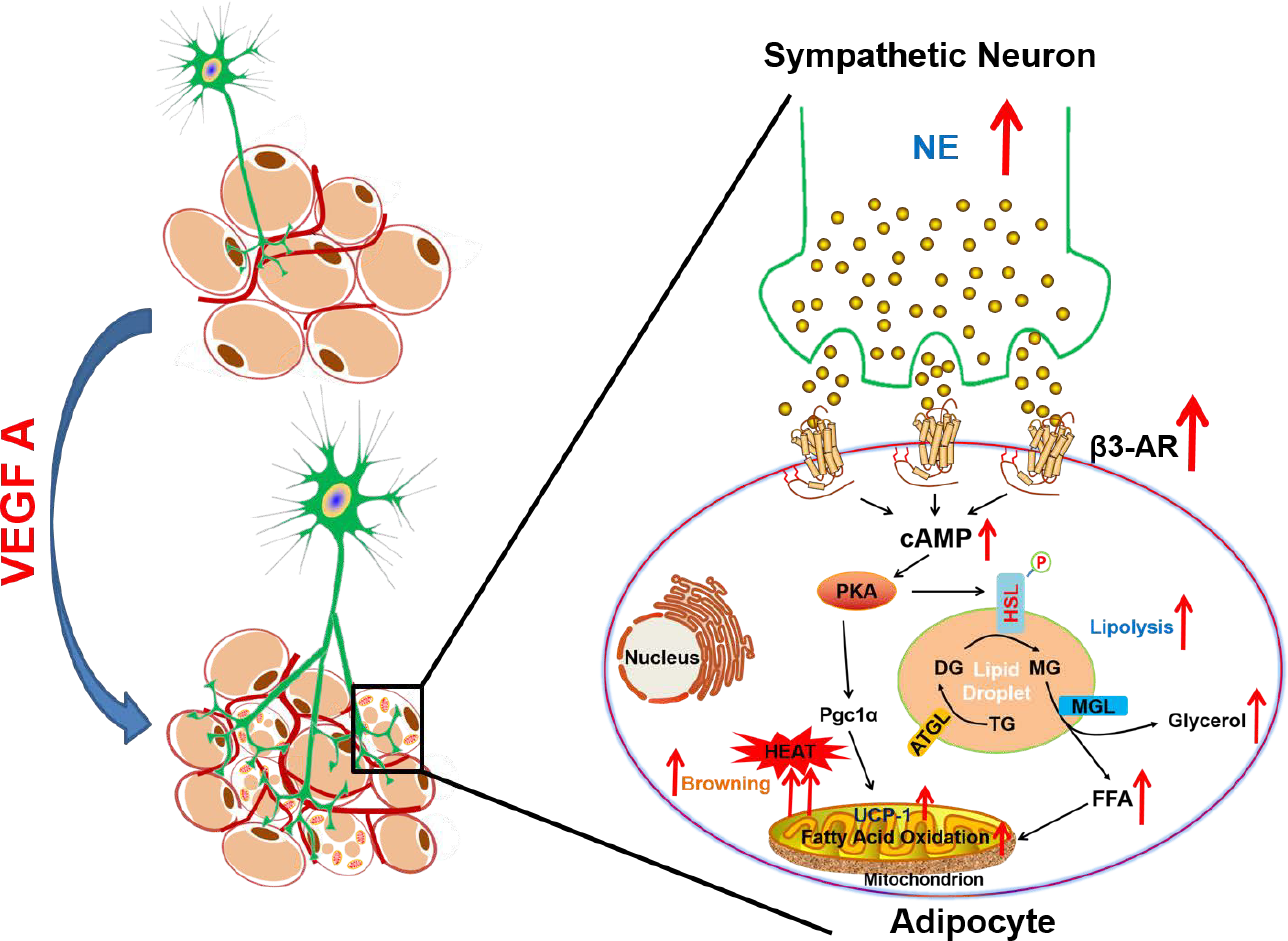
Working model for the functions of VEGF-A in AT. Adipose-derived VEGF-A stimulates activation of sympathetic nervous system which in turn promotes lipolysis and browning, ultimately leading to enhanced energy expenditure.

## Discussion

In response to over-nutritional stress, adipose tissue expands with rapid speed and the blood vessels cannot keep the pace with the expansion [1]. Local hypoxia thus develops. The hypoxic microenvironment in the obese adipose tissue may leads to fibrosis and inflammation which further causes systemic insulin resistance [4, 30, 31]. Previously we and others reported that overexpression of VEGF-A in adipose tissue may counteract with all these adverse effects by inducing angiogenesis and increasing energy expenditure, both of which brings about the metabolic improvement in the diet-induced obese mice [11, 13–16, 19]. Among these reports, the most exciting finding is that in addition to directly activating BAT, VEGF-A stimulates a BAT-like phenotype (browning effect) in sWAT of the transgenic mice [11, 13, 14]. In the current study, we identify the mechanism by which adipocyte-derived VEGF-A promotes lipolysis as well as mitochondrial biogenesis and function in both WAT and BAT. These newly findings, together with the browning effect and BAT activation that we previously reported, clearly explain enhanced energy expenditure in VEGF-A Tg mice. Importantly, the current study further demonstrates that these phenomena are caused by activation of the sympathetic tone locally in the VEGF-A overexpressed adipose tissues.

Of note, while previously we and numerous others have reported metabolically beneficial effects brought about by VEGF-A in adipose tissues, most of these phenomena were observed in the obese mice under HFD feeding for at least 5 weeks [11, 14–16]. Even though the chronic diet-induced obese mouse models are suitable for studies on the anti-obesity effects of VEGF-A, the additional factors that develop during obesity might make it difficult to investigate whether the functions of adipose VEGF-A are direct or indirect. Here, to address the direct effects of VEGF-A in adipose tissue, we took advantage of the Dox-inducible VEGF-A overexpression mouse model and induced the VEGF-A expression only for one week under HFD. Surprisingly, we found that even after the short period of VEGF-A induction, the transgenic mice showed a dramatic phenotype of metabolic improvement including less body weight gains and enhanced energy expenditure. This suggests that adipocyte browning is not an indirect effect of vasculature remodeling but rather a result of direct VEGF-A signaling in adipose tissue.

A growing body of research has demonstrated that nutrient excess leads to mitochondrial dysfunction [32]. For example, obese individuals have been found to have fewer and smaller mitochondria with structural abnormities and reduced oxidative activity. Moreover, the mitochondrial DNA (mtDNA) levels are reduced in the adipose tissue, also supporting a reduction in mitochondria content and function in obesity [33]. These pathological changes dramatically impair the ability of mitochondria to oxidize free fatty acids for use as energy, which causes the adipose tissue to store lipids for fuel instead of burning it. Interestingly, VEGF-A triggered mitochondrial biogenesis and bioenergetics shortly after induction in the transgenic adipose tissues, as demonstrated by upregulated mitochondrial biogenetic and β-oxidation-related genes as well as more mtDNA content, increased mitochondrial density and enhanced OCR. So, how does VEGF-A have the beneficial effects on mitochondria biogenesis and function? One explanation is that in the diet-induced obese adipose tissue, local hypoxia due to the lack of vasculature shapes an unhealthy microenvironment which in turn leads to abnormal mitochondrial functions [1, 14]. Adipose-derived VEGF-A stimulates functional blood vessel formation which hence suppresses the hypoxic conditions by providing adequate oxygen and nutrients [4, 14]. Of high interest in this context is that VEGF-A stimulated massive upregulation of PGC-1α, the primary transcriptional co-factor in the cells which serves as the key driver for mitochondrial biogenesis and function, thereby explaining improved mitochondrial biogenesis and function under HFD challenge. Further studies are needed to identify more detailed molecular mechanism(s) by which VEGF-A stimulates mitochondrial production and function, especially the critical role of nervous innervation in the process.

While sympathetic nerve fibers innervation in adipose tissue is known to be important for the thermogenic function of BAT / beige adipocytes [34], SNS changes during adipose tissue remodeling have not been explored. Recently, the enriched sympathetic nerve fibers as well as neuro-adipose junctions have been observed by using intravital two-photon microscopy [23, 35]. Importantly, *Zeng et al.* further demonstrated that local optogenetic stimulation of the sympathetic fibers induced lipolysis and browning which in turn decreases the fat mass[23, 35]. However, the factors that regulate the sympathetic activation in adipose tissues have remained unclear. **Here, for the first time we showed that VEGF-A stimulates the enhancement of innervation in both sWAT and BAT.** We further demonstrated that the sympathetic pathway plays essential roles in lipolysis and browning in the VEGF-A transgenic mice. Particularly we found that the density of sympathetic nerve fibers were significantly enhanced, as indicated by the IF staining of TH, the marker of sympathetic neurons in both WAT and BAT of the transgenic mice. The local NE levels were increased and the expression of β3-AR was dramatically upregulated in the adipose tissues. As the result, the levels of cAMP were increased and the downstream PKA pathway was activated. The involvement of sympathetic innervation and activation, especially the downstream PKA activation, explains the enhanced lipolysis and browning effects in WAT. Indeed, we found that the levels of phosphorylated HSL by PKA, the critical and rate-limiting lipase in the lipolysis process, were markedly increased and the levels of UCP-1, the marker of the beige cells was massively upregulated on both mRNA and protein levels [14, 19]. Furthermore, we found that the glycerol levels were increased and the smaller as well as multilocular lipid droplets were presented in the sWAT of the VEGF-A transgenic mice. Importantly, both the lipolytic and browning effects were abolished upon β3-adraneceptor antagonist, SR59230A treatment. Collectively, we found that VEGF-A stimulates the increases of the sympathetic drive in WAT. Both lipolysis and browning effects are triggered by the stimulation of β3-AR, which in turn efficiently break down the lipid content for thermogenesis in adipose tissues. Whether lipolysis is necessary for browning remains to be elucidated.

An important question raised from the study is that if VEGF-A triggers the neuronal innervation in adipose tissue, would it do so by inducing angiogenesis alone which in turn upregulates other cytokines to potentiate the neuronal effect or does it directly function on neuronal fibers in the adipose tissues? While previously VEGF-A had been considered to be solely an endothelial mitogen, recent studies demonstrated that VEGF-A has direct actions on neurotrophy, neuronal fiber expansion as well as neuroprotection [36, 37]. Specifically, *Hobson et al.* designed an *in vitro* experiment in a silicone sciatic nerve chamber and found that treatment with VEGF-A dramatically increased nerve axon counts as well as nerve cell proliferation and migration and significantly improved the axon function [38]. This study supports one possibility that in our VEGF-A transgenic adipose tissue, VEGF-A may directly interact with the sympathetic neuronal projection and triggers the fiber proliferation which hence induces the innervation effect. Further studies on novel angiogenic animal models would help to address this question.

In conclusion, our study demonstrates the direct function of VEGF-A on sympathetic activation, which further promotes lipolysis and browning effects in adipose tissue eventually leading to enhanced energy expenditure in the diet-induced obese mice. Since obesity has been considered to be associated with low sympathetic activity [39], our findings thus provide an efficient way to combat/prevent obesity and its associated morbidities.

## Materials and Methods

### Animals

The adipose specific doxycycline (Dox)-inducible VEGF-A transgenic mouse model (VEGF Tg) has been described previously [17]. Briefly, to generate the double transgenic mice, TRE-VEGF Tg mice were crossed with adiponectin promoter-driven-rtTA transgenic mice (Apn-rtTA). The double transgenic mouse (Apn-rtTA-TRE-VEGF-A) expresses VEGF-A in response to Dox treatment. All of these mice are pure C57BL/6J background. All animal experiments were conducted using littermate-controlled male mice, Apn-rtTA Tg mice were used as control mice. All the experiments were started when these mice were 7 weeks old. Mice were housed in cages with 12-dark-light cycle with free access to water and regular chow diet, unless otherwise indicated. The animal study protocols were reviewed and approved by the Animal Welfare Committee of University of Texas Health Science Center at Houston.

### Treatments of Dox-Containing HFD and SR59230A

For all the experiments under HFD plus Dox feeding, the mice were fed with a paste diet containing 60% calories from fat and 30 mg/kg Dox (Bio-Serv, Flemington, NJ) for 7 days; For the treatment of selective β3 adrenoceptor antagonist SR59230A, the VEGF Tg mice were i.p. injected with 1 mg/kg body weight of dissolved SR59230A or with 1% DMSO-PBS placebo every day along with the Dox-containing HFD for 7 days. At day 8, the mice were fasted for 3 h and sacrificed. Tissues and sera were collected for further analyses. SR59230A compound (Sigma-Aldrich, St. Louis, MO) was dissolved in 1% dimethyl sulfoxide (DMSO)-PBS buffer.

### Body Composition and Metabolic Cage Study

Bone-free lean mass, fat mass and body fluids were measured in the non-anesthetized mice by using an Echo3-in-1 NMR (MRI) machine mini Spec (Echo-MRI, Houston, TX). For the indirect calorimetry study, 7 days after HFD plus Dox feeding, the VEGF-A Tg and their littermate control mice were housed individually in TSE metabolic chambers (TSE Systems, Chesterfield, MO) and maintained on a 12 h~12 h dark-light cycle with lights on from 7 AM to 7 PM. All control and VEGF-A Tg mice were fed with HFD plus Dox and water *ad libitum* during the study. Metabolic profiles were recorded continuously for 2 days using TSE metabolic chamber documentation system. The data of the second day were used for analyses.

### Histology

The adipose tissues and the livers were immediately collected after sacrificing the mice and fixed in 10% PBS-buffered formalin (pH 7.4, Fisher Scientific Company, Kalamazoo, MI) for 48 h. Following paraffin embedding, the tissue sections were stained with hematoxylin and eosin (H & E, Sigma-Aldrich) using standard protocol as described previously [4].

For immunohistochemistry (IHC) staining, sections were probed with primary antibody against total phosphorylated PKA substrates (1:1000, Cell Signaling Technologies, Danvers, MA) or against UCP-1 (1:2000, Abcam) followed by biotinylated secondary antibody. Binding of second antibodies were visualized by using DAB chromogen A (Thermo Fisher Scientific, Sugar Land, TX) [4]. Counter-staining was performed with hematoxylin (Sigma-Aldrich). All of the images were acquired with Axio Scope A1 microscope and/or ZEISS microscope software ZEN2 (Carl Zeiss). For immunofluorescence (IF) staining, sections were stained with a primary antibody against COX IV (1:500, Cell Signaling Technologies) followed by a donkey α-rabbit Alexa Fluor 488-conjugated or donkey α-goat Alexa Fluor 647-conjugated secondary antibodies (1.5 μg/mL, Jackson Immuno-Research Laboratories, West Grove, PA). Nuclei were stained with DAPI (1 μg/mL, Vector Laboratories, Burlingame, CA). Images were acquired with Carl Zeiss upright Apotome Axio Imager Z1/ZEN2 Core Imaging software.

### Whole Mount Immuno-Fluorescent Staining and Confocal Microscopy Imaging

The whole mount IF staining for adipose tissues including WAT and BAT has been described previously [24]. Briefly, mice were euthanized by isoflurane and perfused slowly via intracardiac injection with fresh PBS (pH 7.4), followed by removal of the right atrium to release blood. Adipose tissues were excised and were fixed with fresh 1% PBS-buffered paraformaldehyde (PFA, pH 7.4, Electron Microscopy Sciences, Hatfield, PA) for 30 min at room temperature with gentle rocking. After washing with PBS for 3 times, the tissues were subdivided into 0.5-1 cm^3^ sized pieces followed by blocking with 5% BSA in PBS for 30 min at room temperature with gentle rocking. Tissues were then incubated with α-tyrosine hydroxylase (TH) primary antibody (1:1000, Pel Freez Biologicals, Rogers, AR) overnight at 4°C followed by donkey α-rabbit Alexa Fluor 488-conjugated secondary antibody (1.5 μg/mL, Jackson Immuno-Research Laboratories). Nuclei were stained with DAPI (1 μg/mL). Images were acquired with Carl Zeiss upright Apotome Axio Imager Z1/ZEN2 Core Imaging software or confocal Leica TCS SP5 microscope/LAS AF software.

### Quantitative Real-Time PCR (Q-PCR)

Total RNAs was extracted from tissues by TRIzol following the company’s instruction (Invitrogen, La Jolla, CA). The genomic DNAs were removed by DNase I (5 PRIME, Gaithersburg). The total mRNAs were then purified by the RNeasy RNA extraction kit (Qiagen, La Jolla, CA). The quality and quantity of the RNA were determined by 260/280 ratio using Nanodrop 2000 (Thermo Fisher Scientific). cDNAs were obtained by reverse-transcribing 1 μg of total RNAs with iScript Select cDNA Synthesis Kit (Bio-Rad Laboratories, Hercules, CA). Q-PCR reactions were carried out on Bio-Rad CFX96 (Bio-Rad Laboratories). The primer sequences are listed in ***Table 1***. Results were calculated using the 2^−ΔΔCt^ method, normalized by 18s rRNA [4].

**Table 1.**
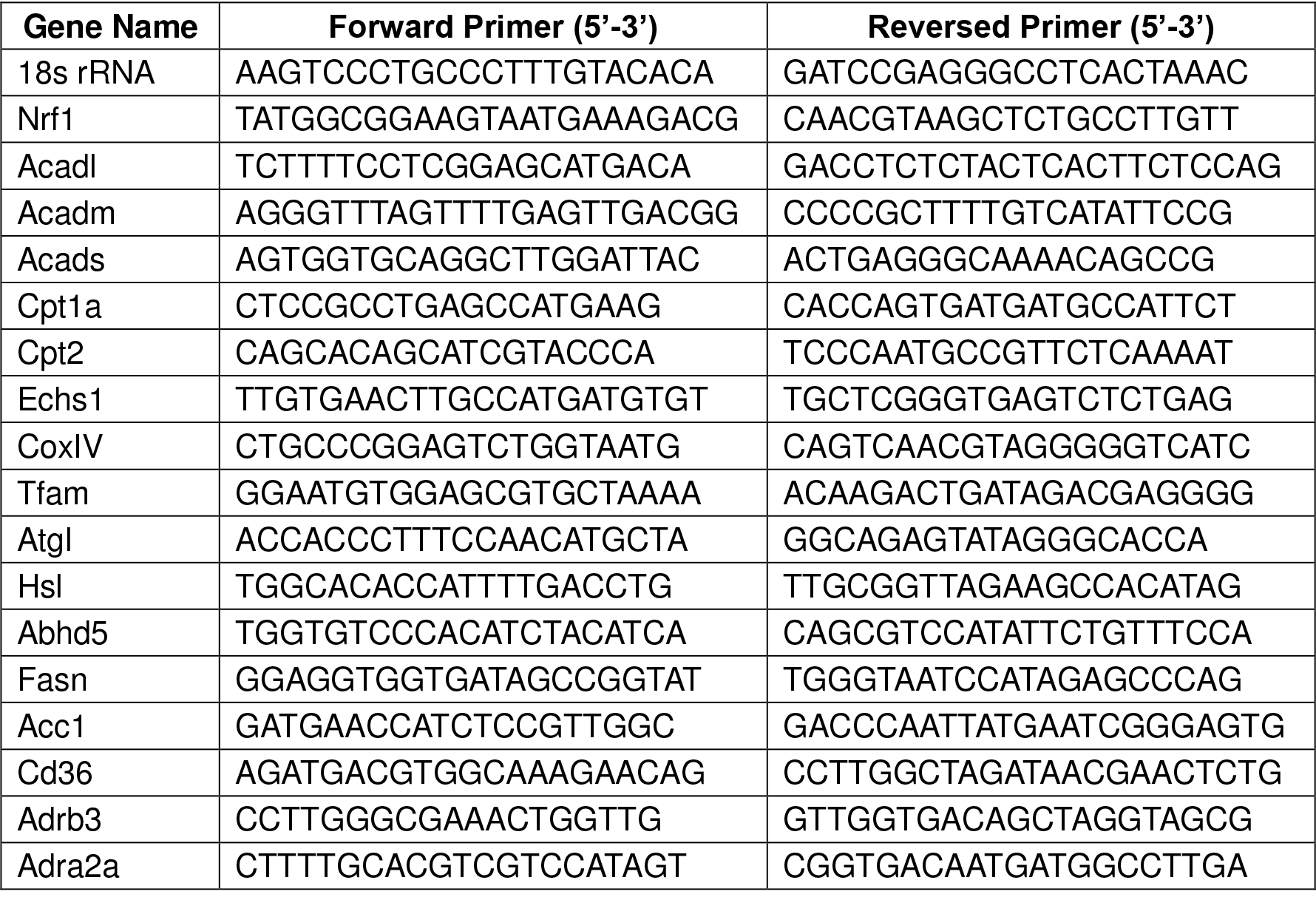
Primer sequences for Q-PCR

### Western Blotting

Total proteins were extracted using cell lysis buffer (25 mM Tris-HCl, 150 mM NaCl,
0. 1% Triton X-100, 0.1% Tween-20, and 0.1% SDS, pH 7.4). The protein concentration was measured by a BCA assay kit (ThermoFisher Scientific). A total 30 μg of cell lysates were separated by 10% or 4-20% Bis-Tris SDS gel (Invitrogen, Carlsbad, CA) and transferred to the PVDF membrane (EMD Millipore, Billerica, MA). For immunoblotting, the primary antibodies for β-actin (1: 2000, Santa Cruz Biotechnology, Dallas, TX), total HSL, phosphorylated HSL (Serine 569/660, PKA targeted sites), or SDHA (1:1000, Cell Signaling Technologies) were used for probing followed by secondary antibodies labeled with infrared dye emitting at 680 nm or 800 nm (Li-Cor Bioscience, Lincoln, NE). The blots were analyzed by Odyssey software (Li-Cor Bioscience). The density of the bands was quantified by ImageJ software (NIH).

### Measurement of NE Levels

The NE levels in serum were detected and quantified in duplicate by an enzyme-linked immunosorbent assay (ELISA) kit (Labor Diagnostika Nord GmbH & Co.KG, Nordhorn, Germany) following the manufacturer’s instruction. In brief, for the NE levels in adipose tissue, WATs were homogenized by sonication in homogenization buffer (1 N HCl, 0.25 M EDTA, and 1M Na_2_S_2_O_5_) and the supernatants were collected by centrifugation at 4°C for NE analyzing after removal of the fat cake at the top. The NE levels were normalized with total amount of tissue proteins.

### Measurement of Glycerol Production

A total 100 mg of WAT were homogenized in 500 μl chilled PBS with proteinase inhibitors (Sigma-Aldrich). Glycerol levels were analyzed by Glycerol Assay Kit (BioAssay Systems, Hayward, CA) following the manufacturer’s instruction. The levels were normalized with total tissue protein levels.

### Measurement of cAMP Levels

cAMP levels in adipose tissue or the serum were detected and quantified in duplicate by an ELISA kit (R & D Systems, Minneapolis, MN) following the manufacturer’s instruction. Briefly, for adipose tissue, WATs were homogenized with a tissue homogenizer in chilled 0.1N HCl at 1:5 ratio (w/v). Supernatants were collected by centrifugation at 4°C and neutralized with 1 N NaOH for analysis.

### Measurement of Oxygen Consumption Rate (OCR) by Seahorse

OCR of WATs was measured in real time by the Seahorse XFe24 Analyzer (Agilent Technologies, Wilmington, DE) as described [40–42]. Briefly, WATs were excised and washed twice with assay medium (DMEM with 25 mM glucose, pH 7.4). Each adipose tissue was sampled using a 2 mm Harris Uni-Core punch (Electron Microscopy Sciences, Hatfield, PA), resulting in 2 mg tissue for each punch. Three tissue punches (totally 6 mg of the tissue) from each fat pad were placed into one well of the XFe24 cell culture microplate (Seahorse Bioscience, North Billerica, MA). Five replicates per fat pad were used for the experiment. The following inhibitors were used at different time windows: 30 μM of oligomycin A, 0.8 μM of carbonyl cyanide-4-(trifluoromethoxy) phenylhydrazone (FCCP), and a mixture of 5 μM of antimycin A and 15 μM of rotenone. All of the inhibitors were obtained from Sigma-Aldrich. The OCR readings were normalized by total amount of DNAs.

### Measurement of Total Lipid Content in Adipose Tissue

A total of 300 mg of WAT excised from the mice was immediately transferred to a 1.5 mL Eppendorf tube with 500 μL of PBS (PH 7.4) at room temperature and was homogenized by MagNA Lyser (Roche Diagnostics, Basel, Switzerland). The homogenized samples were centrifuged at 6000 rpm for 15 min at room temperature to remove the cell debris. The lipid layer was collected and weighed. The amount of lipid content was normalized by total DNAs [31].

### Mitochondrial DNA Assay

A total of 100 mg of WAT was used for nuclear DNA (nuDNA) and mitochondrial DNA (mtDNA) extraction was performed as previously described [43, 44]. The copy numbers of mtDNA were calculated by the ratio of Q-PCR from the mtDNA gene NADH dehydrogenase alpha 1 (NADHA1) to the copy numbers of the nuclear gene lipoprotein lipase (LPL), as previously described [45].

### Statistical Analysis

All data were represented as mean ± SEM. Statistical analysis was performed with GraphPad Prism (GraphPad Software Inc., La Jolla, CA). *Student’s t-test* was applied for statistical analysis. A *p*-value of less than 0.05 was considered to be statistically significant.

## Acknowledgements

We thank Dr. Philipp Scherer at Touchstone Diabetes Center in University of Texas Medical Science Center at Dallas (UTSW) for providing the VEGF-A transgenic animal model. We thank Dr. Zhengmei Mao in the microscopy core of the Institute of Molecular Medicine for assistance on imaging and tissue processing. This study was supported by the National Institute of Health (NIH) grant R01DK109001 (to K.S.) and the Harry E. Bovay, Jr. Foundation.

